# Behavioral and neural mechanisms of face-specific attention during goal-directed visual search

**DOI:** 10.1101/2024.06.24.600413

**Authors:** Jie Zhang, Xiaocang Zhu, Huihui Zhou, Shuo Wang

## Abstract

Goal-directed visual attention is a fundamental cognitive process that enables animals to selectively focus on specific regions of the visual field while filtering out irrelevant information. However, given the domain specificity of social behaviors, it remains unclear whether attention to faces versus non-faces recruits different neurocognitive processes. In this study, we simultaneously recorded activity from temporal and frontal nodes of the attention network while macaques performed a goal-directed visual search task. V4 and inferotemporal (IT) visual category-selective units, selected during cue presentation, discriminated fixations on targets and distractors during the search, but were differentially engaged by face and house targets. V4 and IT category-selective units also encoded fixation transitions and search dynamics. Compared to distractors, fixations on targets reduced spike-LFP coherence within the temporal cortex. Importantly, target-induced desynchronization between the temporal and prefrontal cortices was only evident for face targets, suggesting that attention to faces differentially engaged the prefrontal cortex. We further revealed bidirectional theta influence between the temporal and prefrontal cortices using Granger causality, which was again disproportionate for faces. Finally, we showed that the search became more efficient with increasing target-induced desynchronization. Together, our results suggest domain specificity for attending to faces and an intricate interplay between visual attention and social processing neural networks.

## Introduction

Goal-directed visual attention, a crucial aspect of cognitive processing, reflects the brain’s remarkable ability to prioritize visual information relevant to a specific task or objective. In understanding how animals navigate and interact with their environment, researchers have elucidated neural networks, pathways, and dynamics that govern how the brain selects, processes, and integrates information to achieve desired goals [1–6]. Goal-directed attention involves a complex interplay of various brain regions and circuits, including the prefrontal and temporal cortices [7–12]. These regions work in concert, contributing to the orchestration of attentional resources and forming intricate networks that adaptively modulate sensory processing based on the goals and intentions of the animal [13]. On the other hand, evidence argues for domain specificity in social processing [14, 15]. For example, there is a dedicated face processing system in the macaque inferotemporal (IT) cortex [16, 17]. While faces may inherently attract preferential attention [18], few studies have systematically investigated the detailed behavioral and neurophysiological mechanisms underlying face versus non-face attention. In particular, it remains unclear whether attention to faces recruits different neural substrates and differentially engages the attention neural network.

In this study, we set out to investigate whether visual attention is stimulus-specific and address the debate on whether the processing of faces, a highly social stimulus, in the context of attention is fundamentally different from the processing of non-face stimuli. We employed a free-gaze, goal-directed visual search task using natural face and object stimuli matched for low-level visual properties. Monkeys were trained to detect search targets that belonged to the same category as the cue (but different from the cue), beyond merely matching targets and cues. This allowed for a detailed analysis of attention to faces versus non-faces. We simultaneously recorded from a large number of units from the key brain areas engaged in visual attention and face processing, including V4, IT cortex (TE and TEO), lateral prefrontal cortex (LPFC), and orbitofrontal cortex (OFC). We hypothesize that attention to faces recruits distinct neural processes compared to attention to houses. Furthermore, drawing on prior studies [19, 20], we hypothesize that attention to search targets results in reduced spike-local field potential (LFP) coherence in the lower frequency band. Importantly, we explored whether target-induced desynchronization between the temporal and prefrontal cortices was disproportionate for face targets. Together, by comprehensively analyzing face-specific visual attention through behavior, neuronal firing rate, and functional connectivity, our study contributes to the broader debate on the domain specificity of social processing in the brain (i.e., the notion of the “social brain”).

## Results

### Behavior and eye movement

Two monkeys performed a free-gaze visual search task (**Fig. 1A**) with simultaneous recordings from multiple brain areas (see **Fig. 1C** for detailed characterization of the recording sites). Monkeys were required to fixate on one of the two search targets that belonged to the same category as the cue. Search items were drawn from four categories of natural object images: faces, houses, flowers, and hands, with 40 images per category (**Fig. S1A)**). Importantly, stimuli from different categories were matched in terms of hue and saturation in the HSV color space (**Fig. S1B)**), aspect ratio (**Fig. S1C)**), and luminance (**Fig. S1D)**). This ensured that the findings presented below between faces and houses could not be attributed to low-level features. Both monkeys could well perform this task (accuracy: 91.78%±0.19% for monkey S and 85.23%±0.41% for monkey E). Monkeys had a similar accuracy in finding faces (89.19%±0.31%) versus houses (89.31%±0.33%), although the reaction time (RT), from the onset of the search array to the onset of the last fixation, was faster when searching for faces (**Fig. 1B**; faces: 396.56±1.23 ms, houses: 420.17±1.23 ms; two-tailed Wilcoxon rank-sum test: P = 1.35×10^−166^; Kolmogorov-Smirnov [KS] test: *KS* = 0.081, P = 7.18×10^−228^).

**Fig 1.**
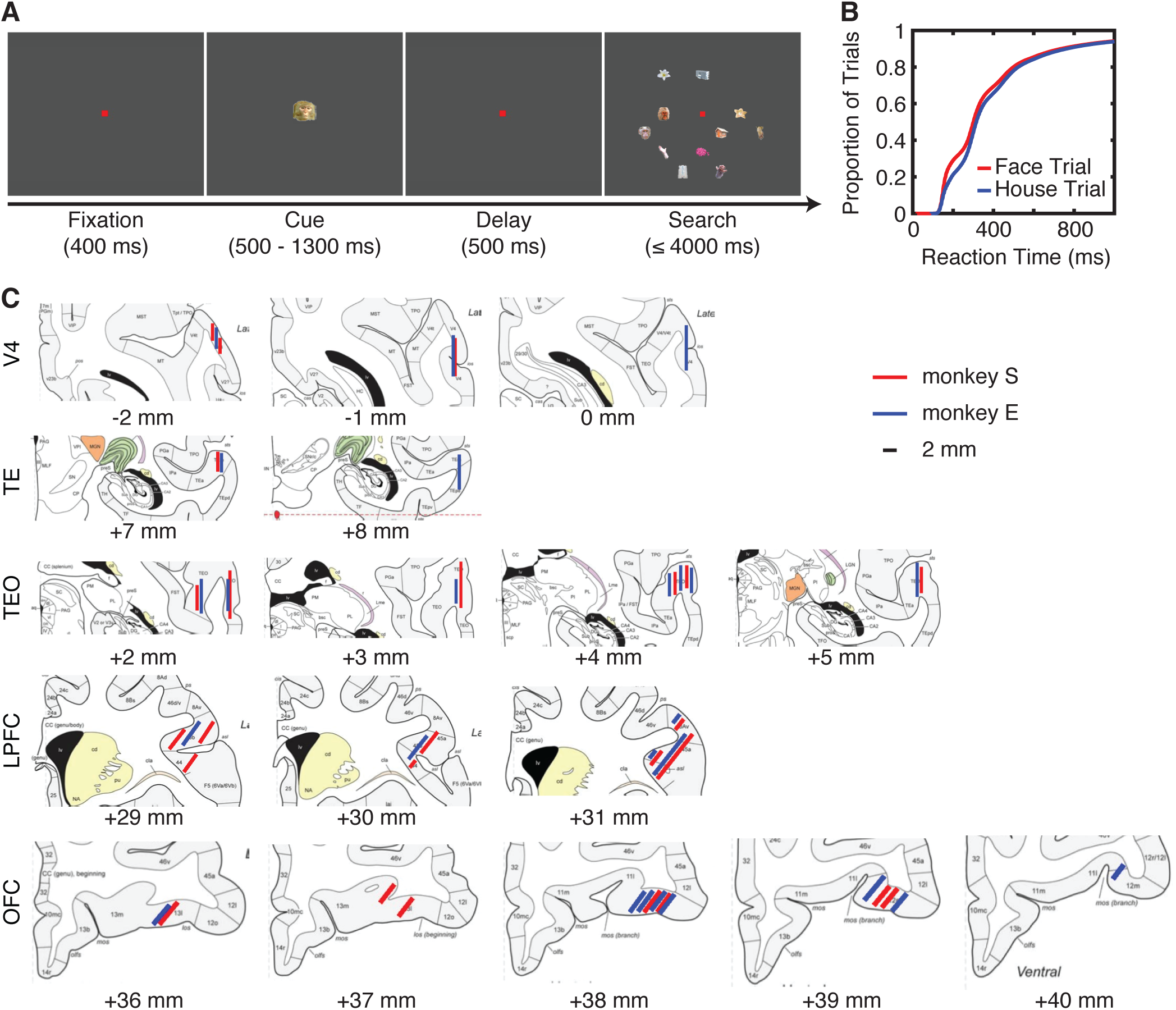
Task, behavior, and recording sites. **(A)** Task. Monkeys initiated the trial by fixating on a central point for 400 ms. A cue was then presented for 500 to 1300 ms. After a delay of 500 ms, the search array with 11 items appeared. Monkeys were required to fixate on one of the two search targets that belonged to the same category as the cue for at least 800 ms to receive a juice reward. **(B)** Cumulative distribution of the reaction time (RT; from the onset of the search array to successful trial completion). **(C)** Recording sites in two monkeys overlaid on the atlas of the rhesus monkey brain in stereotaxic coordinates [21]. The red and blue lines represent the estimated spatial range of recordings in monkey S and monkey E, respectively. The numbers below indicate the rostral (+) or caudal (-) distances of the slices from the infraorbital ridge (Ear Bar Zero).

Monkeys had precise fixations on search items (0.74° to search item center; each search item subtended a visual angle of approximately 2°; **Fig. 2A-C**). We excluded the last fixations on targets from all further analyses where monkeys fixated on the targets for 800 ms to receive juice rewards. We found that fixations on targets was longer than fixations on distractors (**Fig. 2D**; two-way ANOVA [target versus distractor × face versus house]; main effect of target versus distractor: *F*(1,79622) = 9908.87, P < 10^−50^), and fixations on faces was longer than fixations on houses (**Fig. 2D**; main effect of face versus house: *F*(1,79622) = 43.78, P = 3.70×10^−11^; interaction: *F*(1,79622) = 47.51, P = 5.52×10^−12^). Consistent with a longer RT for houses, there were more fixations when monkeys searched for houses compared to faces (**Fig. 2E**; P = 1.75×10^−135^).

**Fig 2.**
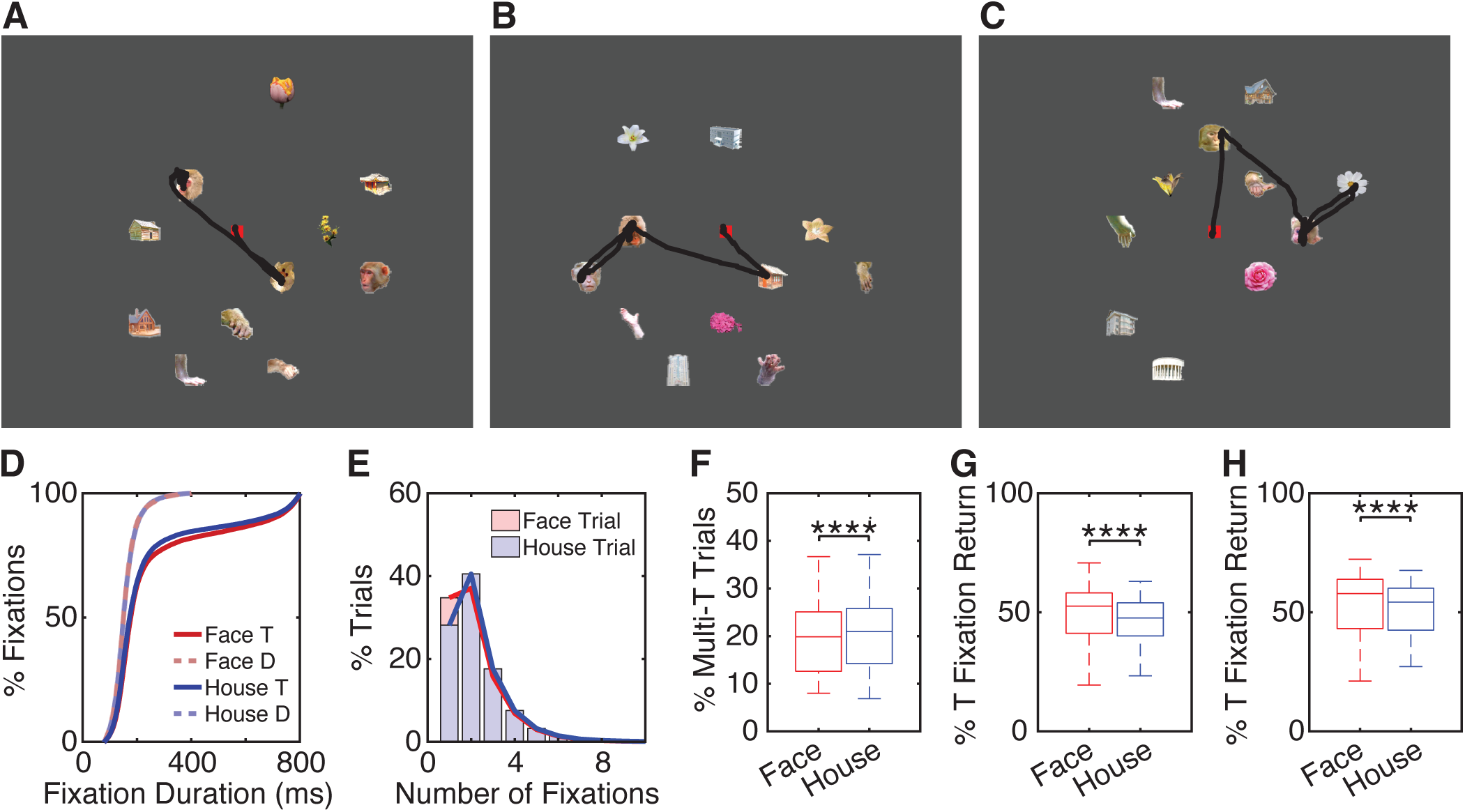
Eye movement during visual search. **(A-C)** Sample search trials. Black dot: raw gaze position. Red square: central fixation (shown during search). **(D)** Cumulative distribution of fixation duration. Trials and fixations were pooled across monkeys, but similar results were derived for each individual monkey. **(E)** Distribution of the number of fixations. **(F)** Percentage of trials with multiple fixations on targets (T). **(G)** Percentage of target (T) fixations that were immediately revisited (with only one item in between; only the same target item was considered). **(H)** Percentage of target (T) fixations that were eventually revisited (with any number of items in between; only the same target item was considered). On each box, the central mark is the median, the edges of the box are the 25th and 75th percentiles, the whiskers extend to the most extreme data points the algorithm considers to be not outliers, and the dots denote the outliers. All box plots in this and subsequent figures follow the same convention. Asterisks indicate a significant difference between conditions using two-tailed Wilcoxon signed-rank test (paired by sessions). ****: P < 0.0001.

In the majority of the trials, monkeys completed the search by fixating on the target when they first spotted it (**Fig. 2A, F**). However, there were trials where monkeys fixated on multiple targets (**Fig. 2B, C, F**), with house targets eliciting more multi-target trials (**Fig. 2F**; P = 1.56×10^−8^). Interestingly, in trials with multiple fixations on targets, a substantial percentage of fixations immediately returned to the same target (**Fig. 2B**; see **Fig. 2C** for a no-return trial; **Fig. 2G**) and eventually returned to the same target (**Fig. 2H**); and notably, face targets had both more immediate (**Fig. 2G**; P = 1.011×10^−8^) and eventual (**Fig. 2H**; P = 2.82×10^−6^) returns. We next explored the neural mechanisms underlying these search behaviors.

### Visual category-selective units signal attention to search targets

We recorded a total of 5070 units from area V4, 5051 units from the IT cortex (including both TE and TEO), 1470 units from the OFC, and 2997 units from the LPFC that had a significant visually evoked response (i.e., the response to the cue or search array was significantly greater than the response to the baseline; Wilcoxon rank-sum test: P < 0.05; see **Fig. 1C** for detailed recording locations). Among these units, 1624 units from V4, 1419 units from the IT cortex, 888 units from the OFC, and 32 units from the LPFC had a focal foveal receptive field (RF; see **Methods**), while 781 units from V4, 268 units from the IT cortex, no units from the OFC, and 514 units from the LPFC had a localized peripheral RF. In this study, we specifically focused on units with a foveal RF to facilitate a comparison between fixations on targets and distractors. Furthermore, given the presence of face-selective regions in the IT cortex [16], we recorded from a wide range of the IT cortex to include units with diverse visual category selectivity (see below; see **Methods**).

We identified visual category-selective units, those distinguishing faces from houses, based on their responses during the cue period (see **Methods**). Our results revealed that 266 units from area V4 (14.01%), 518 units from the IT cortex (34.28%; 10.25% of TEO and 67.02% of TE), 4 units from the LPFC (11.43%), and 640 units from the OFC (71.11%) were *face-preferring* units (**Fig. 3A**; note that both single-unit and multi-unit spikes were included, but we replicated all results with single units only; see below). Similarly, 304 units from area V4 (16.02%), 340 units from the IT cortex (22.50%; 39.68% of TEO and 5.93% of TE), 13 units from the LPFC (37.14%), and 19 units from the OFC (2.11%) were *house-preferring* units (**Fig. 3A**). Importantly, both face-preferring and house-preferring units in V4 (**Fig. 3B**) and IT (**Fig. 3D**) signaled target detection: face-preferring units exhibited a greater response for fixations on face targets compared to fixations on the same faces when they served as distractors (**Fig. 3B, D**). Similarly, house-preferring units exhibited an increased response for fixations on houses when they were targets compared to when they were distractors (**Fig. 3B, D**). Therefore, these results suggest that units in V4 and IT not only signal the categorical membership of the stimuli but also simultaneously convey attention associated with target detection. However, this pattern was not observed for LPFC and OFC category-selective units. Notably, face targets elicited a different attentional effect in both V4 (**Fig. 3C**) and IT (**Fig. 3E**): the response between targets and distractors diverged after approximately 100 ms, and the difference in normalized firing rate was greater for faces than for houses. These results suggest that face and house targets differentially engage these brain areas. Additionally, a separate latency analysis for individual units (see **Methods**) suggested that the attentional effect was earlier for houses than faces in both V4 (**Fig. 3F**; P = 4.40×10^−3^; KS = 0.24, P = 0.010; cf. **Fig. 3C**) and IT (**Fig. 3F**; P = 3.72×10^−7^; KS = 0.26, P = 1.50×10^−5^; cf. the blue versus red top bars indicating a significant difference between targets and distractors shown in **Fig. 3D**).

**Fig 3.**
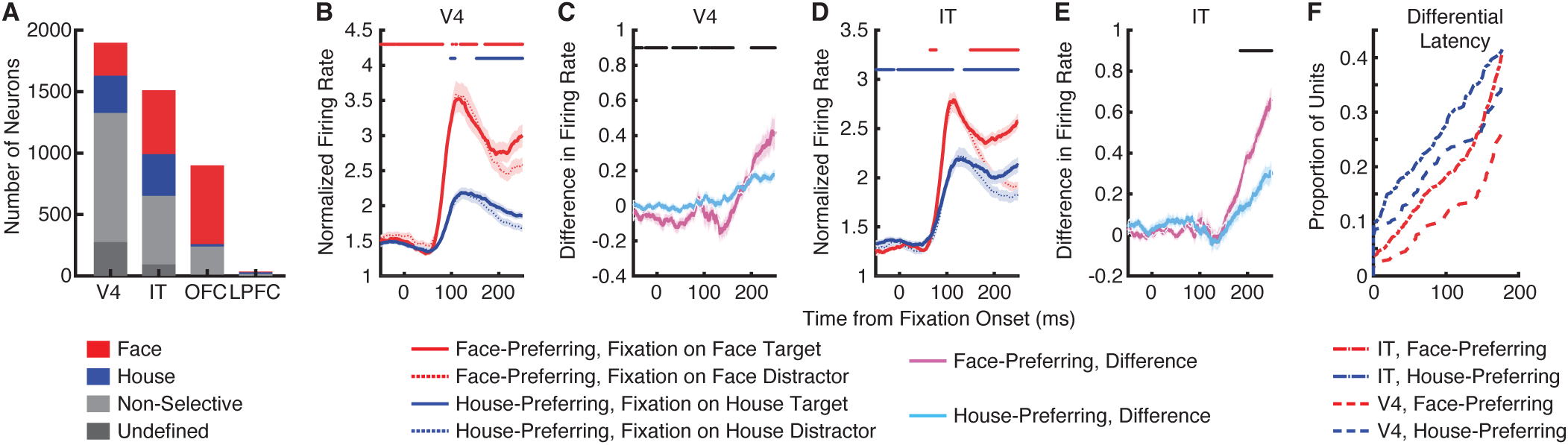
Effect of attention on neuronal firing rate. **(A)** The number of units in each brain area. Red: face-preferring units. Blue: house-preferring units. Light gray: non-category-selective units. Dark gray: undefined units (see **Methods**). **(B, D)** Normalized firing rate. The top bar illustrates the time points with a significant difference between fixations on targets and distractors (two-tailed Wilcoxon rank-sum test, P < 0.05, corrected by FDR [22] for Q < 0.05). Red: fixations on face targets versus fixations on face distractors. Blue: fixations on house targets versus fixations on house distractors. **(C, E)** Difference in normalized firing rate between fixations on targets and fixations on distractors. The top bar illustrates the time points with a significant difference between face-preferring units and house-preferring units (two-tailed Wilcoxon rank-sum test, P < 0.05, corrected by FDR [22] for Q < 0.05). **(B, C)** V4 units. **(D, E)** IT units. Shaded area denotes ±SEM across units. **(F)** Differential latency. The cumulative distribution of differential latencies computed individually for V4 and IT units.

To control for the different levels of response between face-preferring and house-preferring units (**Fig. 3B, D**), we further normalized the difference by the sum of the target and distractor responses, resulting in similar findings (**Fig. S2A, B)**). Additionally, we obtained similar results by normalizing the firing rate by the maximum response of each unit across conditions (**Fig. S2C-F)**). Finally, similar results were obtained using only single units (**Fig. S3**).

### Visual category-selective units signal fixation transitions

Visual search involves complex fixation transitions between search items. We next examined the neural mechanisms underlying these dynamics. First, we observed that category-selective units from the IT cortex (**Fig. 4B**) signaled fixation transitions from search targets. IT category-selective units predicted whether the subsequent fixation was on a target or a distractor, with distractors eliciting a greater response (**Fig. 4A, B**). However, category-selective units did not predict fixation transitions from search distractors (**Fig. 4C, D**; only weakly from the IT cortex). Remarkably, we found that category-selective units could even predict whether fixations would return to the same search target after one fixation (i.e., immediate return; **Fig. 4E, F**) or after a few fixations (**Fig. 4G, H**), particularly for face targets (**Fig. 4E, G**). Together, our results reveal that visual category-selective units can signal fixation transitions and predict search dynamics.

**Fig 4.**
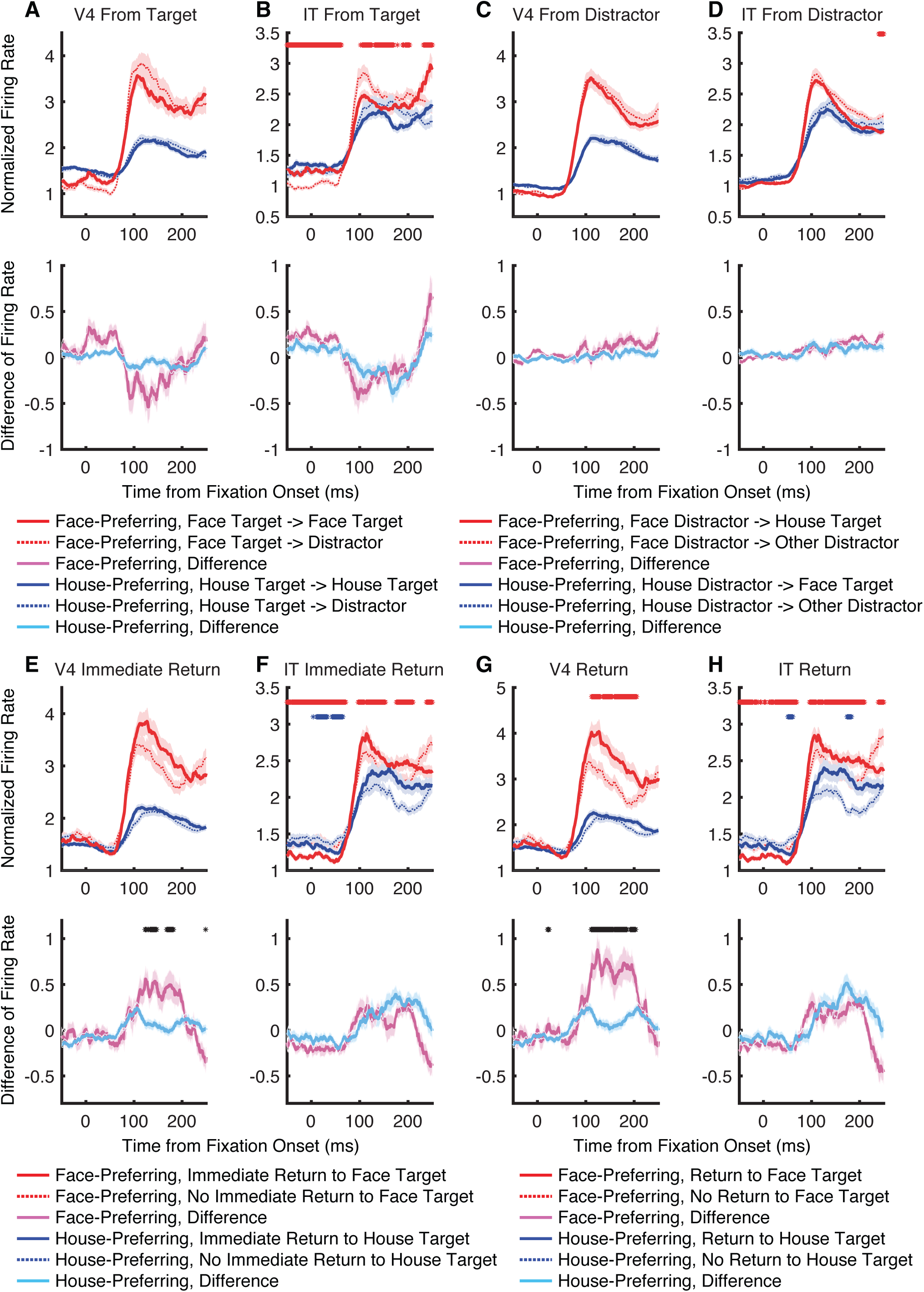
Neural mechanisms of fixation transition. **(A, C, E, G)** V4 units. **(B, D, F, H)** IT units. **(A, B)** Transition from fixation on targets to fixations on targets versus distractors. **(C, D)** Transition from fixation on distractors to fixations on targets versus distractors. **(E, F)** Fixation on targets with (red/blue solid) versus without (red/blue dotted) an immediate return to itself (with only one item in between; only the same target item was considered). **(G, H)** Fixation on targets with (red/blue solid) versus without (red/blue dotted) an eventual return to itself (with any number of items in between; only the same target item was considered). Legend conventions as in Fig. 3.

### Disproportionate target-induced desynchronization for face versus house targets

Does visual search of faces and houses engage the same neural substrates and functional network? To answer this question, we analyzed the coherence between spikes and LFPs recorded simultaneously across brain areas (see **Methods**). We included spikes from V4 and IT (OFC and LPFC were excluded due to having fewer than 20 face-preferring and house-preferring units) and LFPs from all four brain areas. We found that V4 spikes desynchronized with V4 LFPs in the theta frequency band for fixations on targets compared to fixations on distractors, and this was the case for both face targets and house targets (**Fig. 5A**). Such target-induced desynchronization in the theta frequency band was also observed for V4 spike-IT LFP coherence (**Fig. 5B**), IT spike-V4 LFP coherence (**Fig. 5C**), and IT spike-IT LFP coherence (**Fig. 5D**), although house targets elicited a stronger desynchronization between V4 spike and V4 LFP (**Fig. 5A**; two-tailed Wilcoxon rank-sum test between faces and houses: P = 9.51×10^−10^) and between V4 spike and IT LFP (**Fig. 5B**; P = 0.018) whereas face targets elicited a stronger desynchronization between IT spike and IT LFP (**Fig. 5F**; P = 9.64×10^−9^). Importantly, compared to houses, faces elicited significantly greater target-induced desynchronization between V4 spike and OFC LFP (**Fig. 5C**; two-tailed Wilcoxon rank-sum test between faces and houses: P = 7.68×10^−8^), between V4 spike and LPFC LFP (**Fig. 5D**; P = 1.13×10^−4^), and between IT spike and LPFC LFP (**Fig. 5H**; P = 0.0020; note that these results all remained significant with Bonferroni’s correction for 8 comparisons), suggesting that face targets disproportionately engaged the prefrontal cortex compared to house targets.

**Fig 5.**
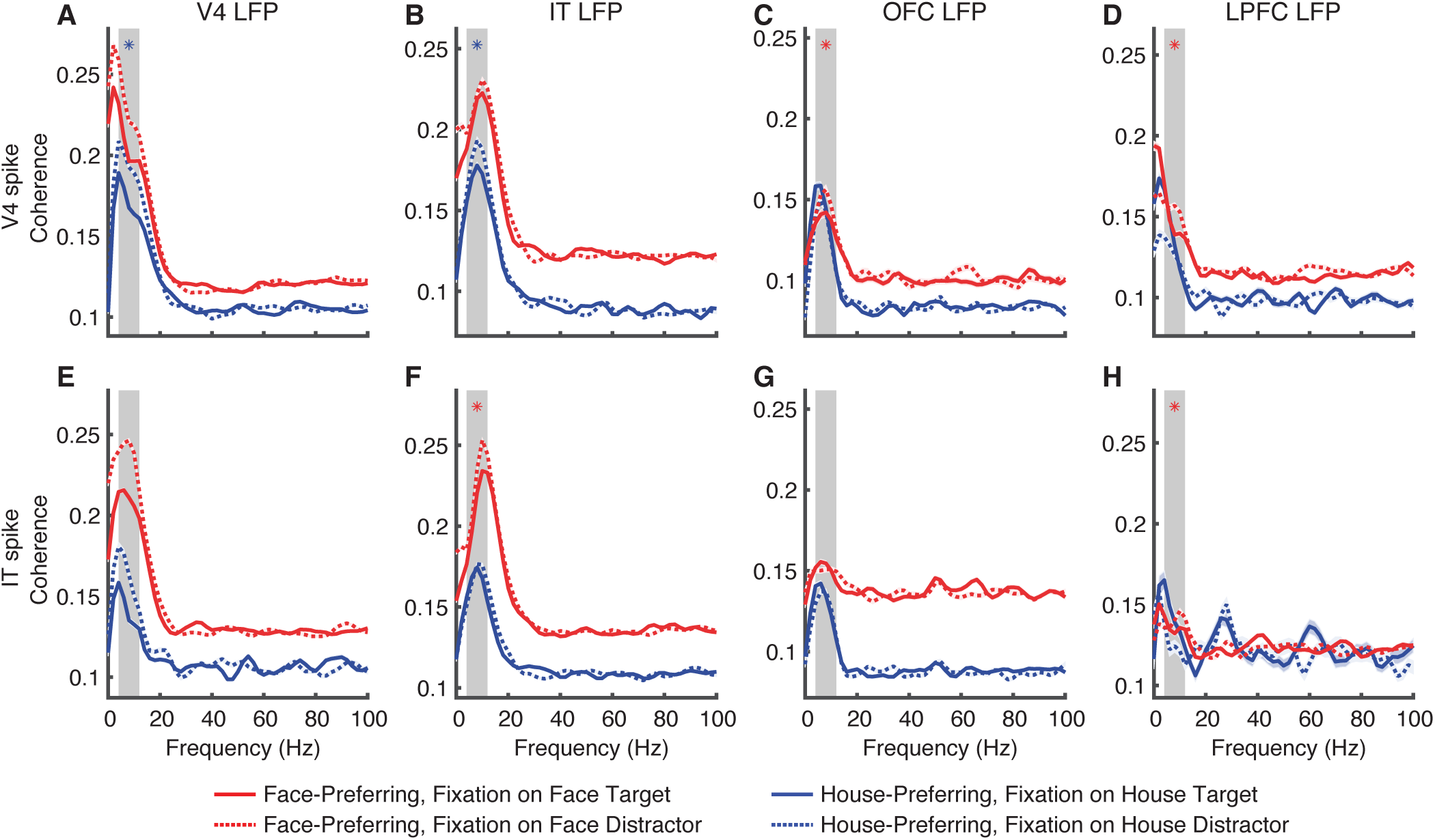
Spike-LFP coherence. **(A)** V4 spike-V4 LFP. **(B)** V4 spike-IT LFP. **(C)** V4 spike-OFC LFP. **(D)** V4 spike-LPFC LFP. **(E)** IT spike-V4 LFP. **(F)** IT spike-IT LFP. **(G)** IT spike-OFC LFP. **(H)** IT spike-LPFC LFP. Red: face fixations. Blue: house fixations. Solid line: fixation on targets. Dotted line: fixation on distractors. Red and blue shaded areas denote ±SEM across spike-LFP pairs. Gray shaded area denotes the theta frequency band (4 - 12 Hz). Asterisks indicate a significant difference in target-induced desynchronization (i.e., the reduction in spike-LFP coherence for fixations on targets compared to fixations on distractors, averaged across the theta frequency band) between face targets and house targets using two-tailed Wilcoxon rank-sum test. Red: face targets > house targets. Blue: house targets > face targets.

It is worth noting that we obtained similar results using only single units (**Fig. S3**). We also obtained similar results when controlling for fixation durations between face targets and house targets as well as between fixations on targets and distractors (for both all units [**Fig. S4A-H**] and single units only [**Fig. S4I-P)**]). Our findings remained robust across various fixation time windows, alignment at saccade onset, different frequency estimations (e.g., choice of multitapers), and units with different types of receptive fields (i.e., focal versus broad). Interestingly, we examined the spike-LFP coherence using the non-preferred stimuli (i.e., houses for face-preferring units and faces for house-preferring units) and found a greater V4-OFC desynchronization for houses in face-preferring units than faces in house-preferring units (for both all units [**Fig. S5A-H)**] and single units only [**Fig. S5I-P)**]), suggesting that target-induced desynchronization with the prefrontal cortex was specific for cell types (i.e., face-preferring units) rather than stimulus types (i.e., faces).

### Directionality of theta influence across brain areas

To investigate the direction of interactions between brain areas, we performed a Granger causality analysis based on spikes and LFPs in the theta frequency band (see **Fig. S6** for analyses across frequencies). We first analyzed the influence of LFPs on spikes (**Fig. 6A-D**; **Fig. S7A-H)**). We found that attention modulated interactions between brain areas when comparing fixations on targets versus distractors, with targets inducing a decrease in Granger causality (i.e., desynchronization). Importantly, such modulation was disproportionate for face versus house targets (**Fig. 6C, D**). Specifically, the influence of V4 LFP on V4 spike (**Fig. S7A)**; two-tailed Wilcoxon rank-sum test between faces and houses: P = 4.01×10^−5^), the influence of OFC LFP on V4 spike (**Fig. 6A**; **Fig. S7C)**; P = 5.88×10^−11^), and the influence of LPFC LFP on IT spike (**Fig. 6B**; **Fig. S7H)**; P = 0.0018) were more strongly modulated by face targets, whereas the influence of IT LFP on V4 spike (**Fig. S7B)**; P = 0.0050) and the influence of IT LFP on IT spike (**Fig. S7F)**; P = 0.0011) were more strongly modulated by house targets.

**Fig 6.**
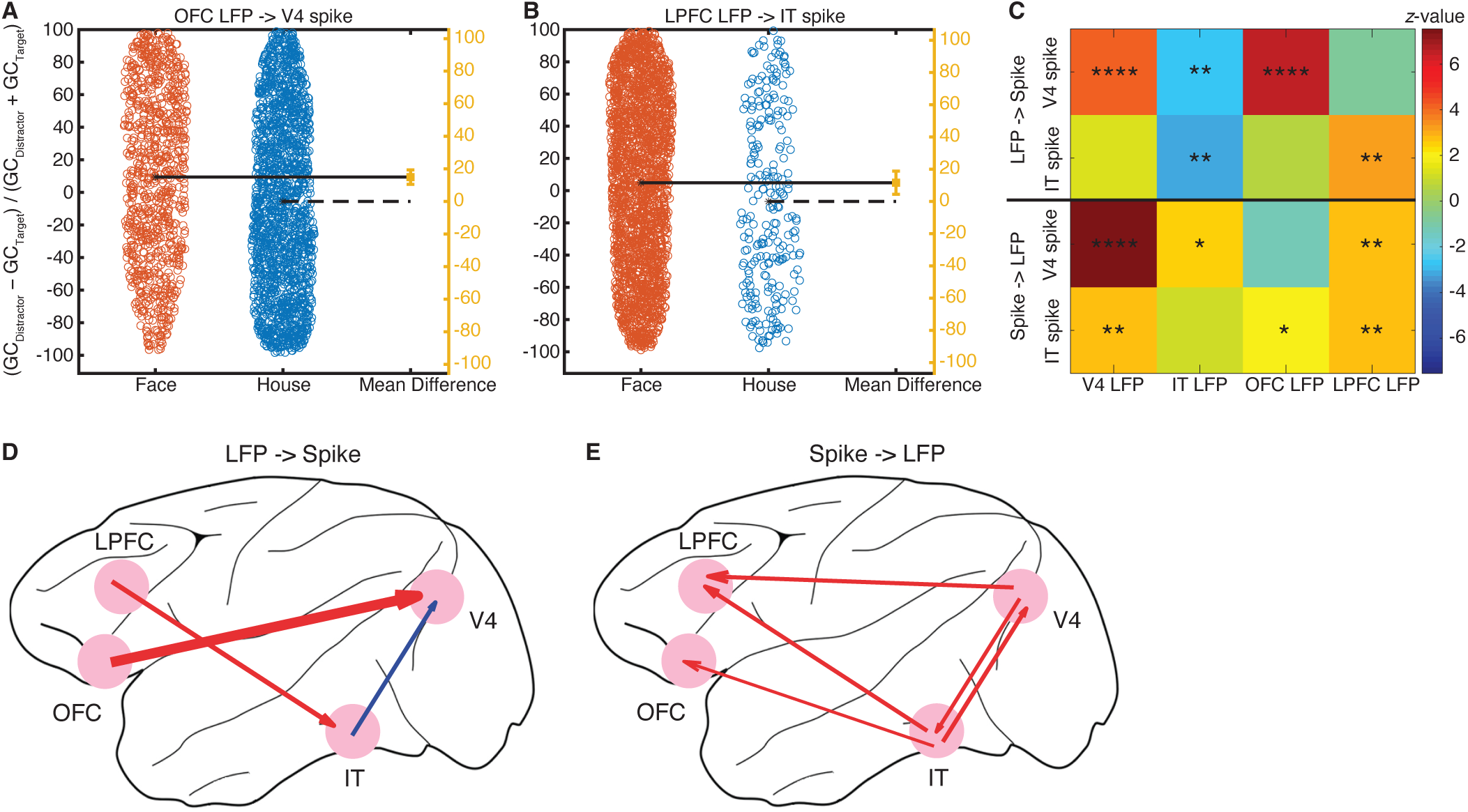
Granger causality (GC). **(A, B)** Compared to house targets, face targets induced a stronger reduction in Granger causality between the temporal and prefrontal cortices ([GC_Distractor_ − GC_Target_] / [GC_Distractor_ + GC_Target_]). Each circle represents a spike-LFP pair. The mean of the house targets corresponds to the zero effect size and the mean of the face targets corresponds to the value of the effect size on the effect size axis (yellow). The vertical error bar in yellow indicates the actual mean-difference effect size value and the confidence intervals. **(A)** OFC LFP influence on V4 spike. **(B)** LPFC LFP influence on IT spike. **(C)** Summary of Granger causality for each spike-LFP pair. Color coding shows the *z*-values of the Wilcoxon rank-rum test (face − house). Asterisks indicate a significant difference between face targets and house targets using two-tailed Wilcoxon rank-sum test. *: P < 0.05, **: P < 0.01, ***: P < 0.001, and ****: P < 0.0001. **(D, E)** Significant differences in target-induced reduction of cross-area Granger causality between face and house targets. The thickness of the arrow is proportional to the *z*-value of the Wilcoxon rank-rum test. Red: face > house. Blue: house > face. **(D)** LFP influence on spike. **(E)** Spike influence on LFP.

We also analyzed the influence of spikes on LFPs (**Fig. 6C, E**; **Fig. S7I-P)**). Again, we found that attention modulated interactions between brain areas, and such modulation was disproportionate for face and house targets (**Fig. 6C, E**). Notably, the influence of V4 spike on V4 LFP (**Fig. S7I)**; P = 4.62×10^−14^), the influence of V4 spike on IT LFP (**Fig. S7J)**; P = 0.015), the influence of V4 spike on LPFC LFP (**Fig. S7L)**; P = 0.0077), the influence of IT spike on V4 LFP (**Fig. S7M)**; P = 0.0055), the influence of IT spike on OFC LFP (**Fig. S7O)**; P = 0.047), and the influence of IT spike on LPFC LFP (**Fig. S7P)**; P = 0.0074) were all more strongly modulated by face targets.

Lastly, we calculated spike-spike Granger causality (see **Methods**) between V4 and IT (note that the OFC and LPFC were excluded from this analysis because they had fewer than 20 face-preferring and house-preferring units) and found that the influence of V4 spike on IT spike was more strongly modulated by house targets (P = 0.0034). However, we did not observe a significant difference between face and house targets for the influence of IT spike on V4 spike (P = 0.53).

Together, our results reveal bidirectional influences between LFPs and spikes in the temporal and prefrontal cortices. In particular, attention modulation of these influences is more pronounced for face targets compared to house targets, with face targets showing a disproportionate engagement of the prefrontal cortex (see **Fig. 6C-E** for a summary).

### Relationship between behavior, firing rate, and spike-LFP coherence

Finally, we investigated the relationship between search efficiency (indexed by RT and the number of fixations), attentional effect (indexed by the difference in firing rate for targets versus distractors), and target-induced desynchronization (indexed by the reduction in spike-LFP coherence for targets compared to distractors). Specifically, we found that RT correlated with the target-induced reduction in IT spike-V4 LFP coherence (**Fig. 7A**), especially for house targets (Pearson correlation; face: *r* = −0.11, P = 0.016; house: *r* = −0.57, P = 2.33×10^−8^; note that only house targets remained significant with Bonferroni correction for 8 comparisons). On the other hand, RT correlated with the target-induced reduction in V4 spike-V4 LFP coherence only for face targets and not house targets (**Fig. 7B**; face: *r* = −0.24, P = 1.18×10^−4^; house: *r* = −0.013, P = 0.83). Furthermore, RT correlated with the target-induced reduction in IT spike-IT LFP coherence for both face and house targets (**Fig. 7C**; face: *r* = −0.12, P = 0.0051; house: *r* = −0.20, P = 2.27×10^−4^). Similarly, the number of fixations per trial correlated with the target-induced reduction in IT spike-V4 LFP coherence for house targets (**Fig. 7D**; face: *r* = −0.11, P = 0.019; house: *r* = −0.58, P = 9.83×10^−9^), V4 spike-V4 LFP coherence for face targets (**Fig. 7E**; face: *r* = −0.24, P = 8.22×10^−5^; house: *r* = 0.012, P = 0.84), and IT spike-IT LFP coherence for both face and house targets (**Fig. 7F**; face: *r* = −0.13, P = 0.005; house: *r* = −0.18, P = 0.0011). However, we did not observe significant correlations between search efficiency and attentional effects (all Ps > 0.05), nor between attentional effects and target-induced desynchronization (all Ps > 0.05).

**Fig 7.**
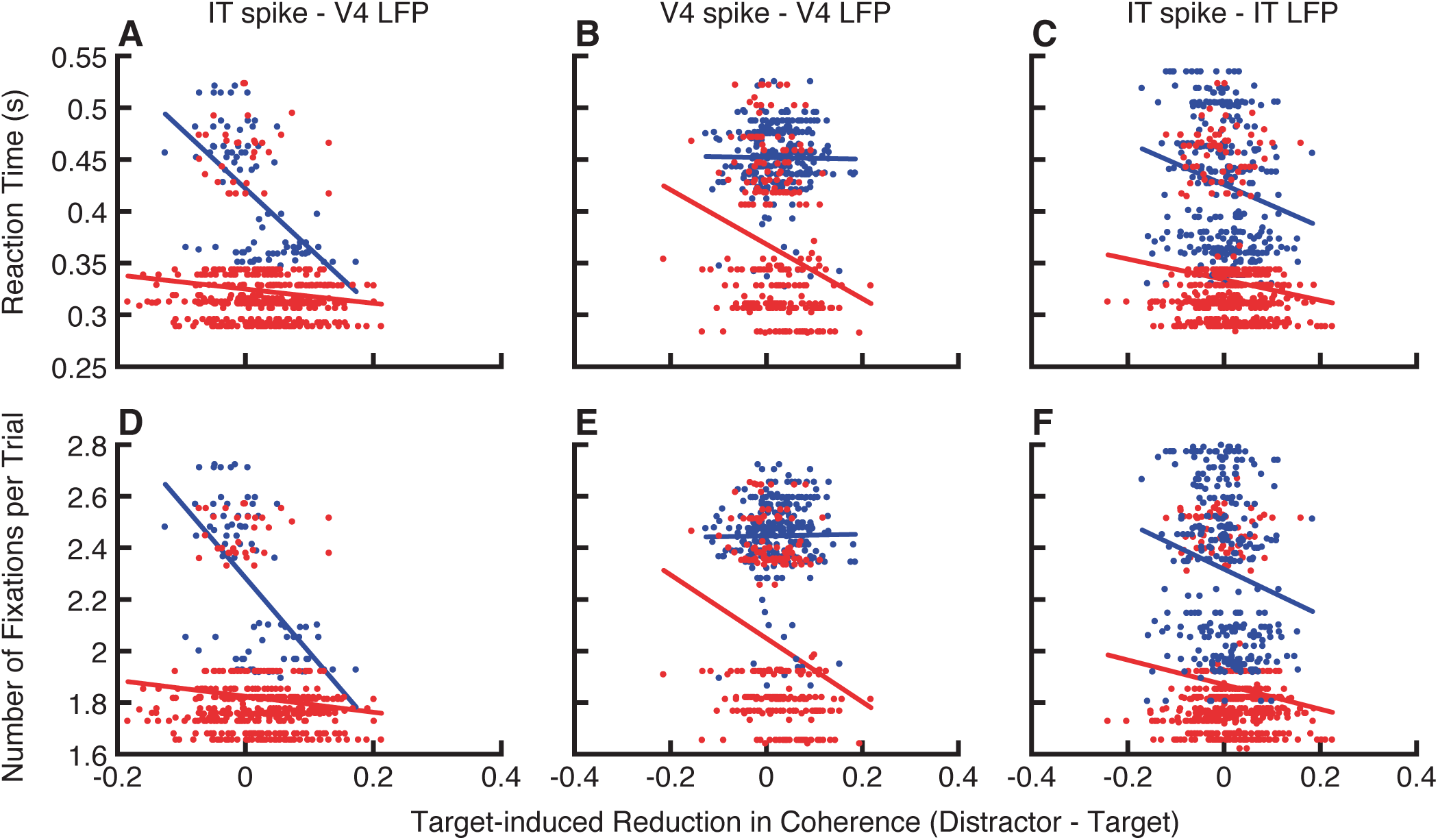
Relationship between behavior and spike-LFP coherence. **(A)** Correlation between reaction time (RT) and target-induced reduction in IT spike-V4 LFP coherence (distractor − target). **(B)** Correlation between RT and target-induced reduction in V4 spike-V4 LFP coherence. **(C)** Correlation between RT and target-induced reduction in IT spike-IT LFP coherence. **(D)** Correlation between the number of fixations and target-induced reduction in IT spike-V4 LFP coherence. **(E)** Correlation between the number of fixations and target-induced reduction in V4 spike-V4 LFP coherence. **(F)** Correlation between the number of fixations and target-induced reduction in IT spike-IT LFP coherence. Each dot represents a unit, and the lines represent the linear fit. Red: face targets. Blue: house targets.

Together, our results suggest that target-induced desynchronization within the temporal cortex can explain search efficiency: search becomes more efficient (reduced RT and number of fixations) with increasing desynchronization. Notably, faces and non-faces exhibit different patterns of correlation between neural responses and search behavior, further indicating that different neural processes are involved in searching for faces.

## Discussion

In this study, we demonstrated that visual category-selective units in V4 and IT, selected during cue presentation, distinguished between fixations on targets and distractors during visual search. We also elucidated the neural mechanisms underlying fixation transitions and search dynamics, particularly noting differences between face and house targets. Furthermore, we observed reduced spike-LFP coherence in V4 and IT with increased attention to search targets, and such target-induced desynchronization between temporal and prefrontal cortices was only evident for face targets. Finally, we revealed directional theta influences between the temporal and prefrontal cortices under attention modulation, which was again disproportionate for face targets. Together, our results suggest domain specificity in searching for faces, as well as an intricate interplay between visual attention and face processing. Notably, we demonstrated not only local neural computations by showing the attentional effect for neurons within a brain area but also information flow across brain areas using functional connectivity. These neural mechanisms were further linked to behaviors.

Our study represents one of the first examinations of feature attention in foveal units during visual search, complementing the common view of distributed feature attentional effects [8, 23] and providing a more comprehensive understanding of the distribution of attentional effects across the entire visual field. Notably, foveal units facilitate the examination of target-induced attentional effects and the comparison between face and house targets because they exhibit stronger category selectivity and more focused receptive fields than peripheral units in V4 and IT [24]. Foveal attentional enhancements may aid in maintaining eye fixation on the current target, potentially reducing the likelihood of making saccades towards peripheral items during search [24]. Interestingly, we also demonstrated that category-selective units predicted fixation transitions and search dynamics. Importantly, all these aspects differed between face and house targets. While our results could not be simply attributed to low-level visual features (**Fig. S1**), future studies are needed to elucidate the underlying differences between face and non-face stimuli. This includes investigating whether our findings could be explained by factors such as familiarity, motivation, value, and attractiveness.

We observed desynchronization for attended stimuli (fixations on search targets) compared to the same stimuli that were unattended (fixations on distractors) in the theta frequency band in both V4 and IT, consistent with prior studies showing desynchronization for attended stimuli in V4 in a similar frequency band [19, 20]. Desynchronization has been observed for both feature-based attention (distinguishing between target and distractor in the peripheral receptive field) and saccade selection (directing attention into [attention in] or out of [attention out] the peripheral receptive field), and it was the case for V4 spike-V4 LFP coherence, V4 spike-frontal eye field (FEF) LFP coherence, and FEF spike-V4 LFP coherence [20]. Importantly, we showed that even when the search task remained the same and the accuracy remained similar, the processing for face targets disproportionately engaged the prefrontal cortex compared to house targets. Our results thus suggest a context modulation of the attention neural network based on stimulus category, contrasting with the invariant V4 response to different reward/motivation contexts [25]. This result may also explain the advantages of searching for faces, as well as the disproportionate deficits in attention to faces observed in individuals with autism spectrum disorder during visual search [26].

The neural responses observed when fixating on or saccading to search targets versus distractors are commonly interpreted as indicative of attentional processes [8, 11]. However, similar to many other studies, we cannot entirely rule out the influence of reward expectation on these neural responses. It is plausible that both template matching and reward expectation contribute to the observed attentional effects. This dual role underscores the complexity of interpreting neural responses in search tasks and suggests that attentional and reward-related processes are intricately linked. Further research is needed to disentangle these contributions and clarify the mechanisms underlying these neural responses. Furthermore, while the reward was the same for face and non-face trials, faces were detected faster than houses (**Fig. 1B**), indicating that detecting faces requires less mental effort. However, reaction time, which indicates stimulus difficulty and mental effort, correlated only with spike-LFP coherence within the temporal cortex (i.e., V4 and IT), but not with attentional effects in firing rate (**Fig. 7**). Therefore, the differences in neural response could not be simply explained by differences in behavior. In particular, behavioral differences could not explain the differential engagement of the prefrontal cortex for face targets (**Fig. 5**) or the differential attentional effect for faces (**Fig. 3**).

Simultaneous neural recordings across brain areas have delineated the roles of different nodes within the attention neural network. For example, it has been shown that synchrony between LPFC and parietal areas is stronger in lower frequencies during top-down attention and in higher frequencies during bottom-up attention [9]. Paired neuron recordings in FEF and IT reveal that spatial selection precedes object identification during visual search [27]. Top-down attention originates in the prefrontal and parietal cortices and influences the sensory temporal cortex both through direct descending projections (e.g., from FEF to V4) and through a backward cascade (e.g., from the prefrontal cortex, particularly LPFC, to IT to V4) [13]. Our present study not only supports the backward cascade but also demonstrates a differential backward cascade based on stimulus type. The disproportionate engagement of the prefrontal cortex may result from different interactions between excitatory pyramidal neurons and inhibitory interneurons, which are central to the mechanisms supporting normalization and the generation of synchronous oscillations [13]. Future studies will be needed to investigate whether attention to faces versus non-faces differentially recruits other nodes of the attention network, such as the FEF [28]. It also remains an interesting question to explore whether the backward progression of attentional effects in the ventral stream [29] is similar for faces versus non-faces.

One critical question regarding the notion of the “social brain” is whether any of the neural networks exhibit specialization for processing social information. Earlier notions propose that social processing in primates is subserved by a specific brain system [14], which is supported by neurons dedicated to face processing in the IT cortex, OFC, and amygdala [16]. More recent views tie subsets of the social processing structures together into functional networks that serve particular components of social cognition [15]. Consistent with our previous findings that single neurons in the human prefrontal cortex [30] and medial temporal lobe [31] encode visual search targets and demonstrate category-selective responses to faces [31], our present study provides a network view integrating visual attention and category selectivity during visual search. In particular, our findings that processing social information differentially engages the prefrontal cortex support domain-specific neural processing of social stimuli [14, 15, 18]. Together, our comprehensive analyses of behavior, neuronal firing rate, and functional connectivity collectively advance our understanding of the neural mechanisms underlying visual social attention and provide strong support for the domain specificity of social processing and the notion of the “social brain”. Our study contributes to this debate by providing a detailed analysis of how faces (as social stimuli) and non-face stimuli are processed differently in terms of visual attention.

## Acknowledgements

This research was supported by the NSF (BCS-1945230), NIH (R01MH129426), and AFOSR (FA9550-21-1-0088). The funders had no role in study design, data collection and analysis, decision to publish, or preparation of the manuscript.

## Author Contributions

J.Z. and H.Z. designed research. J.Z. and X.Z. performed experiments. J.Z. and S.W. analyzed data. J.Z., H.Z., and S.W. wrote the paper. All authors discussed the results and contributed toward the manuscript.

## Competing Interests Statement

The authors declare no conflict of interest.

## Methods

### Subjects and recording sites

Two male rhesus macaques, weighing 12 and 15 kg, were used in the study. All experiments were performed with the approval of the Institutional Animal Care and Use Committee of Shenzhen Institutes of Advanced Technology, Chinese Academy of Sciences (No. SIAT-IRB-160223-NS-ZHH-A0187-003).

The monkeys were implanted under aseptic conditions with a post to fix the head and recording chambers over areas V4, inferotemporal (IT) cortex (including both TE and TEO), lateral prefrontal cortex (LPFC), and orbitofrontal cortex (OFC; see **Fig. 1C** for details). The localization of the chambers was based on MRI scans obtained before surgery. Notably, given the presence of face-selective regions in the IT cortex [16], we recorded neural activity from a wide range of the IT cortex to include units with diverse visual category selectivity. The recording sites were determined based on stereotaxic coordinates relative to Ear Bar Zero (EBZ) in the atlas of the rhesus monkey brain and the known regions of face patches [32]. Recordings spanned from the central IT cortex, encompassing the area between the anterior middle temporal sulcus (AMTS) and the posterior middle temporal sulcus (PMTS). TE recordings were concentrated from +7 to +8 mm rostral to EBZ, near the posterior edge of the AMTS, while TEO recordings were concentrated from +2 to +5 mm rostral to EBZ, near the anterior edge of the PMTS (**Fig. 1C**). Most face-preferring units were located from +7 to +8 mm rostral to EBZ, likely within the ML face patch in TE. In contrast, most house-preferring units were found from +2 to +5 mm rostral to EBZ, specifically within the TEO area.

### Tasks and stimuli

Monkeys were trained to perform a free-gaze visual search task. A central fixation was presented for 400 ms, followed by a cue lasting 500 to 1300 ms. After a delay of 500 ms, the search array was on. The search array contained 11 items, including two targets, randomly selected from a total of 20 predefined locations. Monkeys were required to find either one of the two targets within 4000 ms and maintain fixation on the target for 800 ms to receive a juice reward. No constraints were placed on their search behavior to allow animals to perform the search naturally. Before the onset of the search array, monkeys were required to maintain a central fixation. The two target stimuli belonged to the same category as the cue stimulus, though they were distinct images. We utilized four categories of stimuli—face, house, flower, and hand—each comprising 40 images. The cue stimulus was randomly selected from the house or face stimuli with equal probability. The remaining 9 stimuli in the search array were drawn from the other three categories. Each stimulus subtended an area of approximately 2° × 2°, with the hue, saturation in the HSV color space, aspect ratio, and luminance of these images matched across categories. The 20 locations, covering the visual field of eccentricities from 5° to 11°, included 18 locations located symmetrically in the left and right visual field, with 9 on each side, and 2 locations on the vertical middle line.

A visually guided saccade task was employed to map the peripheral receptive fields (RFs) of recorded units. Following a 400-ms central fixation, a stimulus randomly appeared in one of the 20 locations, and monkeys were required to make a saccade to the stimulus within 500 ms and maintain fixation on it for 300 ms to receive a reward.

Behavioral experiments were conducted using the MonkeyLogic software (University of Chicago, IL), which presented the stimuli, monitored eye movements, and triggered the delivery of the reward.

### Electrophysiology

Single-unit and multi-unit spikes were recorded from V4, IT, LPFC, and OFC using 24- or 32-contact electrodes (V-Probe or S-Probe, Plexon Inc, Dallas, USA) in a 128-channel Cerebus System (Blackrock Microsystems, Salt Lake City, UT, USA). In most sessions, we recorded activities in two of the areas simultaneously. Neural signals were filtered between 250 Hz and 5 kHz, amplified, and digitized at 30 kHz to obtain spike data. The recording locations in V4, IT, LPFC, and OFC were verified with MRI. Eye movements were recorded using an infrared eye-tracking system (iViewX Hi-Speed, SensoMotoric Instruments (SMI), Teltow, Germany) at a sampling rate of 500 Hz.

### Data analysis: spike rate

Measurements of neural activity were obtained from spike density functions, which were generated by convolving the time of action potentials with a function that projects activity forward in time (Growth = 1 ms, Decay = 20 ms) and approximates an EPSP [33]. The spike rate of each unit was normalized by the mean baseline firing rate during the fixation spot preceding the cue.

### Data analysis: receptive field

The visual response to the cue and the search array in the free-gaze visual search task was assessed by comparing the firing rate during the post-stimulus period (50 to 200 ms after cue/array onset) to the corresponding baseline (−150 to 0 ms relative to cue/array onset) using a Wilcoxon rank-sum test. Based on these responses, we classified units into three categories of RFs:

i. Units with a focal foveal RF: These units responded solely to the cue in the foveal region (P < 0.05) but not to the search array that included items in the periphery (P > 0.05).
ii. Units with a broad foveal RF: These units responded to both the cue and the search array.
iii. Units with a peripheral RF: These units only responded to the search array (P < 0.05) but not to the cue (P > 0.05). The RFs of these units were additionally mapped based on their activities in the visually guided saccade task.

Units not classified into one of the three categories were excluded from further analysis. In this study, our focus was on units with a foveal RF (categories (i) and (ii)), comparing fixations on targets versus distractors.

### Data analysis: category selectivity

We determined the category selectivity of each unit by comparing the response to face cues versus house cues in a time window of 50 to 200 ms after cue onset (Wilcoxon rank-sum test, P < 0.05). We further imposed a second criterion using a selectivity index similar to indices employed in previous IT studies [34, 35]. For each unit with a foveal RF, the response to face stimuli (*R*_face_) or house stimuli (*R*_house_) was calculated using the visual search task by subtracting the mean baseline activity (−150 to 0 ms relative to the onset of the cue) from the mean response to the face or house cue (50 to 200 ms after the onset of the cue). The selectivity index (SI) was then defined as (*R*_face_ − *R*_house_) / (*R*_face_ + *R*_house_). SI was set to 1 when *R*_face_ > 0 and *R*_house_ < 0, and to −1 when *R*_face_ < 0 and *R*_house_ > 0. Face-preferring units were required to have an *R*_face_ at least 130% of *R*_house_ (i.e., the corresponding SI was greater than 0.13). Similarly, house-preferring units were required to have an *R*_house_ at least 130% of *R*_face_ (i.e., the corresponding SI was smaller than −0.13). Units were labeled as non-category-selective if the response to face cues versus house cues was not significantly different (P > 0.05). The remaining units that did not fit into any of the aforementioned types were classified as undefined units (i.e., there was a significant difference but did not meet the second criterion).

### Data analysis: attentional effect

We calculated the attentional effect as the difference in firing rate between the same stimuli when they served as targets versus distractors. The latency of attentional effect at the population level was determined based on the mean response of each unit using a sliding window method. If a significant difference (Wilcoxon signed-rank test, P < 0.05) was found successively for 35 ms between the target and distractor responses, the first time point of the 35 ms window was defined as the starting point of the attentional effect. To test whether a latency difference at the population level was significant, we used a two-sided permutation test with 1000 runs, as described in our previous study [11]. The latency of the attentional effect for each unit was defined as the first 20-ms bin out of the twelve successive bins that had a significantly greater response for targets than distractors (one-tailed Wilcoxon signed-rank test: P < 0.05).

### Data analysis: spike-LFP coherence

We implemented the spike-LFP coherence analysis using the Chronux toolbox (www.chronux.org) in MATLAB. We used a single Hanning taper across frequencies, but we derived similar results using multitaper methods for higher frequencies (> 25 Hz) [36]. Coherence between two signals, *x* and *y*, was calculated using the following formula:

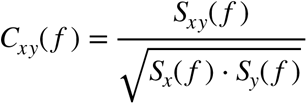

where *S_x_*(*f*) and *S_y_*(*f*) denote the auto-spectra and *S_xy_*(*f*) represents the cross-spectrum of the two signals *x* and *y*. Auto-spectra and cross-spectra were averaged across trials before coherence calculation. We used a 200-ms time window for each fixation (0 to 200 ms relative to fixation onset). Notably, we used an equal number of fixations and an equal number of spikes for fixations on targets and distractors to calculate coherence for a given pair of recording sites, thus eliminating bias from different sample sizes. To avoid spikes contributing to the LFP recorded on the same electrode, we utilized signals from two different electrodes to calculate coherence. Furthermore, we used the same LFP signals for both face targets and house targets, without selecting LFPs based on their category selectivity, attention selectivity, or the selectivity of the associated units (e.g., the LFP signals could come from contacts with both face-preferring units and house-preferring units). Lastly, although we used category-selective units with preferred stimuli for analysis, non-category-selective units showed consistent results.

### Data analysis: Granger causality

We utilized the open-source MATLAB toolbox “SpikeFieldGrangerCausality” [37] for frequency-domain Granger causality analysis between spikes and LFP. Causality was calculated during the same period as in coherence analysis between spikes and LFP across various brain areas. The power spectrum for the spiking process was estimated with the multitaper method and then frequency-domain Granger causality from LFP *xi* to spike *dNj* was calculated as follows:

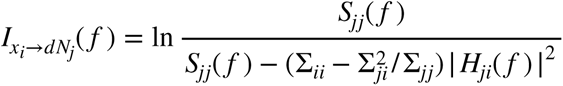

where *S_jj_*(*f*) is the auto-spectra of spike point process at frequency *f*, *H* is the transfer matrix, and *Σ* is the noise covariance matrix. The influence in the opposite direction (from spike *dN_i_* to LFP *x_j_*) can be analyzed in a similar way.

We utilized the open-source MATLAB toolbox “gcpp” [38] to assess Granger causality between spikes of multiple neurons during the same period as described above. A potential causal relationship from neuron(s) *j* to neuron *i* is assessed using the log-likelihood ratio, which is given by:

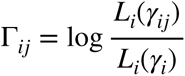

where *L_i_*(*γ_i_*) is the likelihood of producing a particular set of spike trains of neuron *i* that is calculated using all the covariates of neuron *i*, and *L_i_*(*γ_i_^j^*) is the likelihood of neuron *i* excluding the spiking history of neuron(s) *j*. The Granger causality measure from neuron(s) *j* to neuron *i* is calculated as:

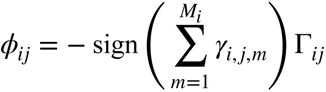

where the sign of 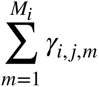 represents an averaged influence of the spiking history of neuron(s) *j* on neuron *i*, which distinguishes excitatory and inhibitory influences. The population average over the absolute value of *Φ_ij_* was used for comparison across brain areas.

---

**Fig S1.**
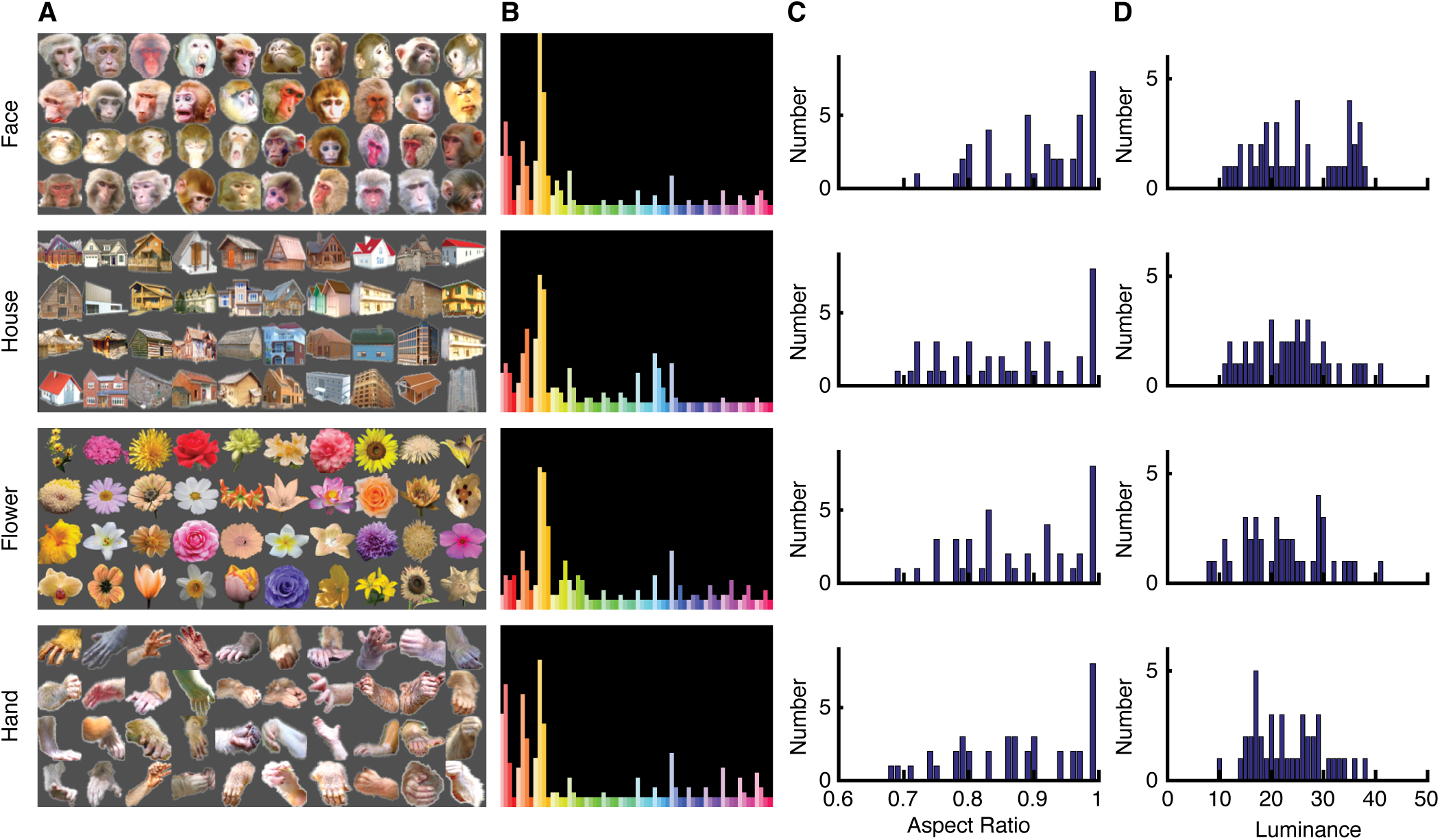
Characterization of the stimuli. **(A)** Four categories of visual objects (40 images per category) were used for neural recordings. **(B)** The stimuli from different categories did not exhibit significant differences in pixel-wise hue and saturation (χ^2^-test: P > 0.05). **(C)** The stimuli from different categories did not exhibit significant differences in aspect ratio (P > 0.05). **(D)** The stimuli from different categories did not exhibit significant differences in luminance (P > 0.05). Shown are histograms for each stimulus category.

**Fig S2.**
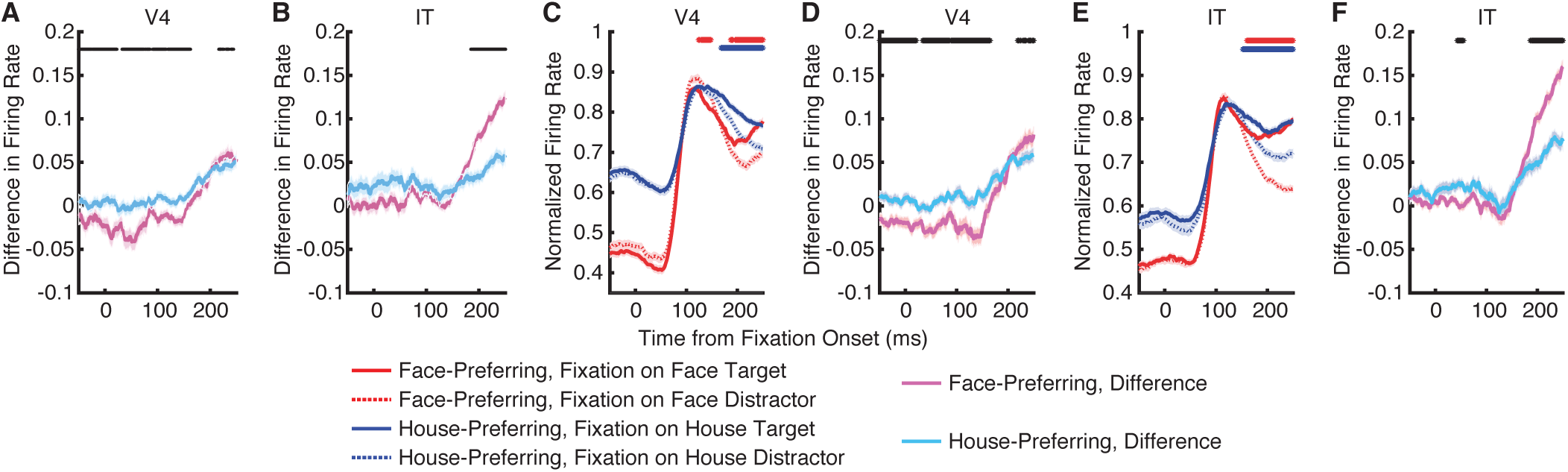
Effect of attention on neuronal firing rate. **(A, B, D, F)** Difference in normalized firing rate between fixations on targets and fixations on distractors. **(C, E)** Normalized firing rate. **(A, B)** The difference in baseline-normalized firing rate was further normalized by the sum of target and distractor response at each time point. **(C-F)** The firing rate was normalized by the maximum response of each unit across conditions. Legend conventions as in Fig. 3.

**Fig S3.**
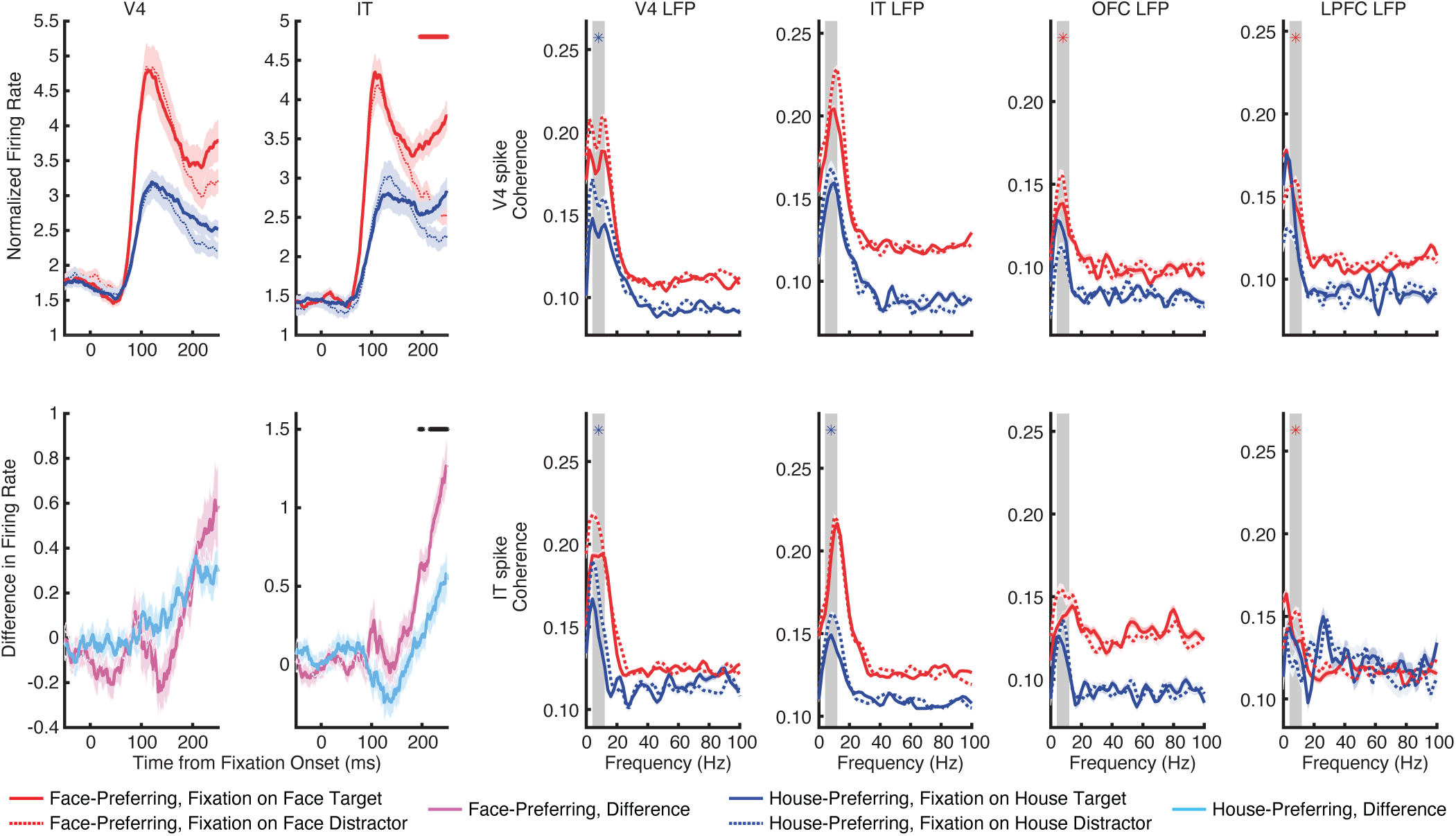
Effect of attention on neuronal firing rate and spike-LFP coherence for single units only. Legend conventions as in Fig. 3 and Fig. 5.

**Fig S4.**
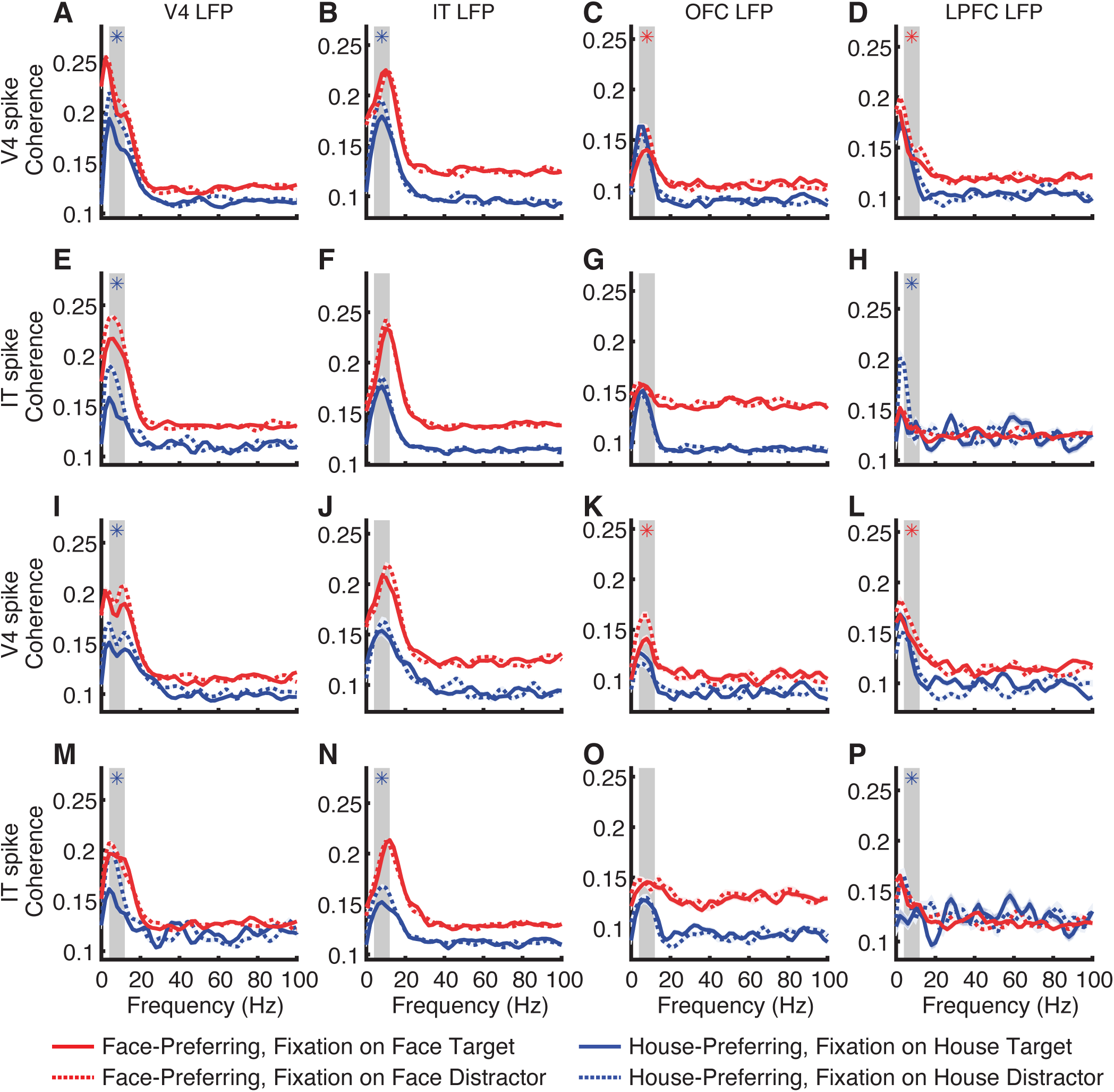
Spike-LFP coherence for fixations with equal durations. **(A-H)** All units. **(I-P)** Single units only. Legend conventions as in Fig. 5.

**Fig S5.**
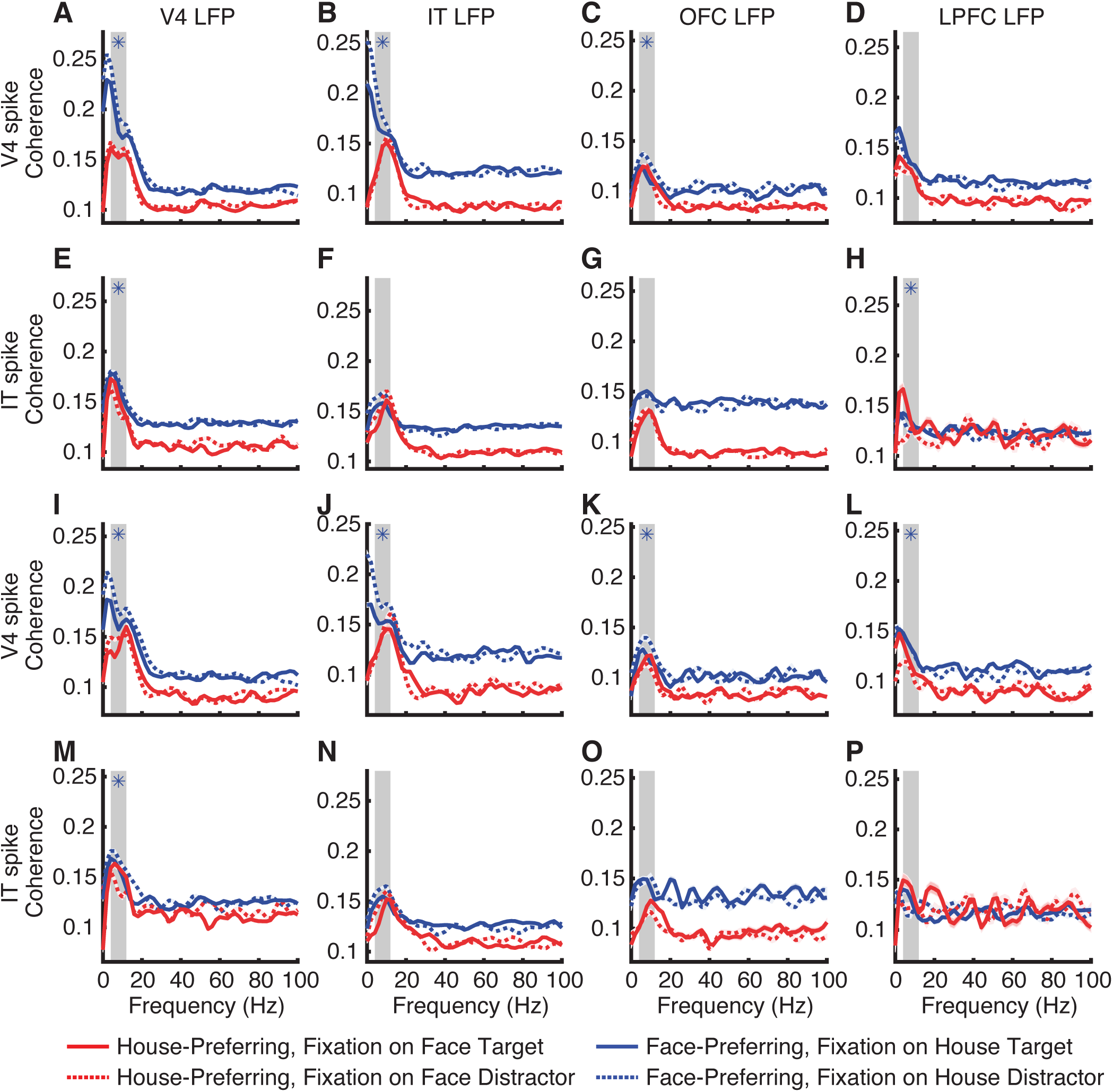
Spike-LFP coherence for non-preferred stimuli. **(A-H)** All units. **(I-P)** Single units only. Legend conventions as in Fig. 5.

**Fig S6.**
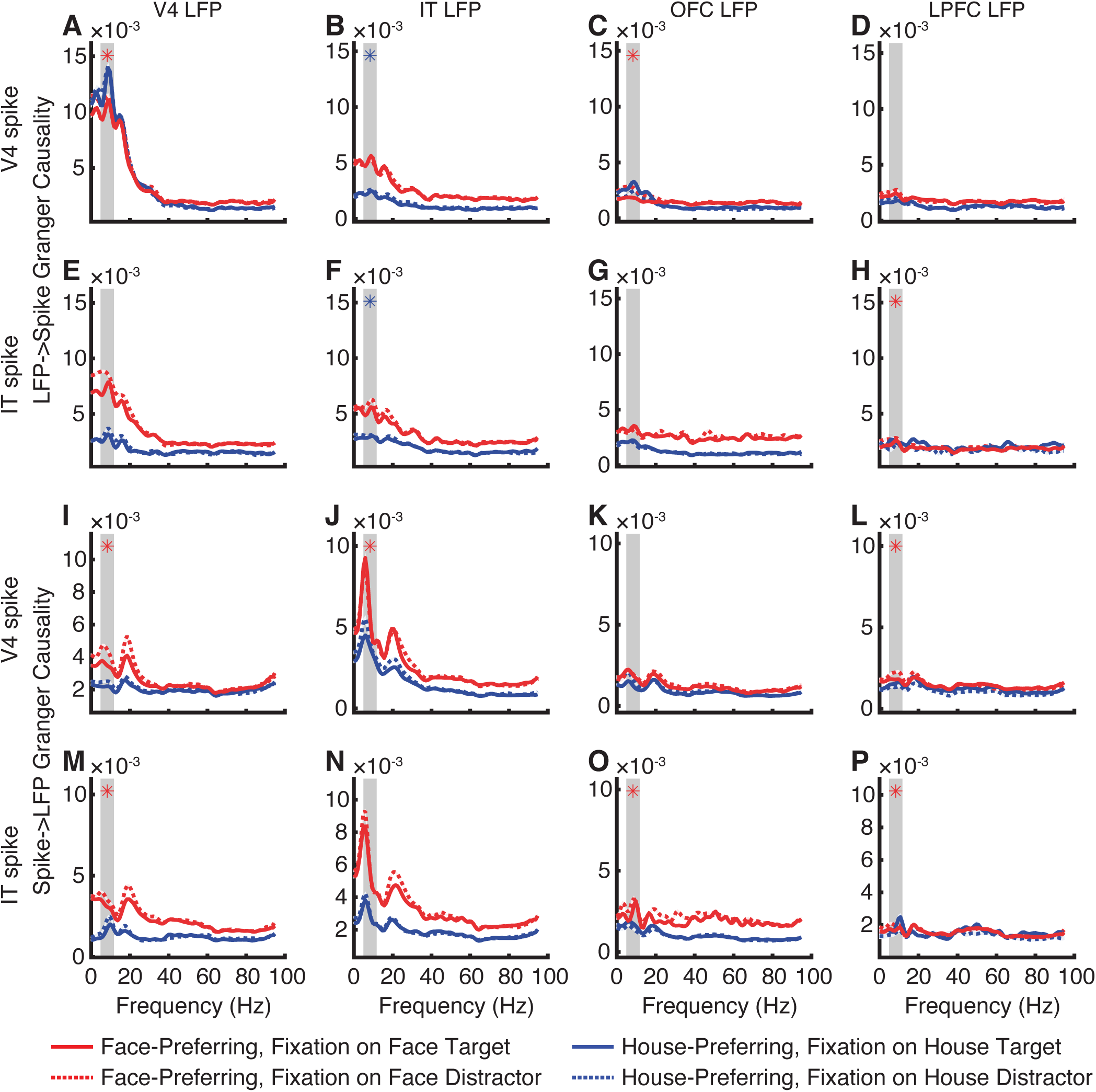
Granger causality. **(A)** V4 LFP influence on V4 spike. **(B)** IT LFP influence on V4 spike. **(C)** OFC LFP influence on V4 spike. **(D)** LPFC LFP influence on V4 spike. **(E)** V4 LFP influence on IT spike. **(F)** IT LFP influence on IT spike. **(G)** OFC LFP influence on IT spike. **(H)** LPFC LFP influence on IT spike. **(I)** V4 spike influence on V4 LFP. **(J)** V4 spike influence on IT LFP. **(K)** V4 spike influence on OFC LFP. **(L)** V4 spike influence on LPFC LFP. **(M)** IT spike influence on V4 LFP. **(N)** IT spike influence on IT LFP. **(O)** IT spike influence on OFC LFP. **(P)** IT spike influence on LPFC LFP. Legend conventions as in Fig. 5.

**Fig S7.**
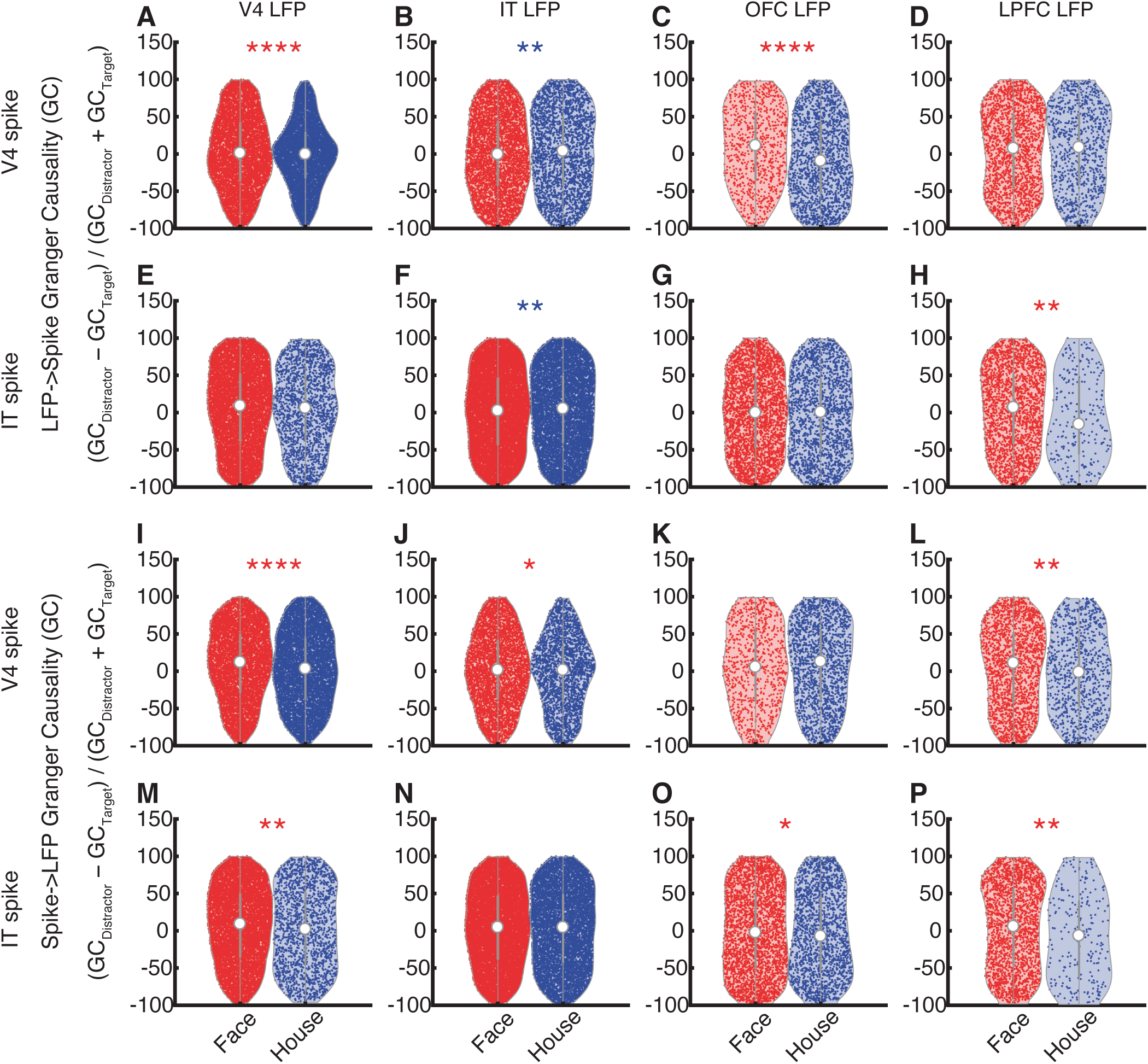
Granger causality (GC). **(A)** V4 LFP influence on V4 spike. **(B)** IT LFP influence on V4 spike. **(C)** OFC LFP influence on V4 spike. **(D)** LPFC LFP influence on V4 spike. **(E)** V4 LFP influence on IT spike. **(F)** IT LFP influence on IT spike. **(G)** OFC LFP influence on IT spike. **(H)** LPFC LFP influence on IT spike. **(I)** V4 spike influence on V4 LFP. **(J)** V4 spike influence on IT LFP. **(K)** V4 spike influence on OFC LFP. **(L)** V4 spike influence on LPFC LFP. **(M)** IT spike influence on V4 LFP. **(N)** IT spike influence on IT LFP. **(O)** IT spike influence on OFC LFP. **(P)** IT spike influence on LPFC LFP. In the violin plots, the white dot represents the median, the thick gray bar in the center represents the interquartile range, the thin gray line represents the rest of the distribution, except for points that are determined to be outliers using a method that is a function of the interquartile range. On each side of the gray line is a kernel density estimation to show the distribution shape of the data. Asterisks indicate a significant difference between face targets and house targets using two-tailed Wilcoxon rank-sum test. *: P < 0.05, **: P < 0.01, ***: P < 0.001, and ****: P < 0.0001. Red: face targets > house targets. Blue: house targets > face targets.

## Notes

### Competing Interest Statement

The authors have declared no competing interest.

